# Neuroanatomical diversity of corpus callosum and brain volume in the Autism Brain Imaging Data Exchange (Abide) project

**DOI:** 10.1101/002691

**Authors:** Aline Lefebvre, Anita Beggiato, Thomas Bourgeron, Roberto Toro

**Author notes:** Corresponding author: Roberto Toro, PhD, Institut Pasteur, 25, rue du Docteur Roux, 75015, Paris, France.

## Abstract

The corpus callosum – the main pathway for long-distance inter-hemispheric integration in the human brain – has been frequently reported to be smaller among autistic patients compared with non-autistic controls. We conducted a meta-analysis of the literature which suggested a statistically significant difference. However, the studies included were heavily underpowered: on average only 20% power to detect differences of 0.3 standard deviations, which makes it difficult to establish the reality of such a difference. We therefore studied the size of the corpus callosum among 694 subjects (328 patients, 366 controls) from the Abide cohort. Despite having achieved 99% power to detect statistically significant differences of 0.3 standard deviations, we did not observe any. To better understand the neuroanatomical diversity of the corpus callosum, and the possible reasons for the previous findings, we analysed the relationship between its size, the size of the brain, intracranial volume and intelligence scores. The corpus callosum appeared to scale non-linearly with brain size, with large brains having a proportionally smaller corpus callosum. Additionally, intelligence scores correlated with brain volume among controls but the correlation was significantly weaker among patients. We used simulations to determine to which extent these two effects could lead to artefactual differences in corpus callosum size within populations. We observed that, were there a difference in brain volume between cases and controls, normalising corpus callosum size by brain volume would not eliminate the brain volume effect, but adding brain volume as a covariate in a linear model would. Finally, we observed that because of the weaker correlation of intelligence scores and brain volume among patients, matching populations by intelligence scores could result in a bias towards including more patients with large brain volumes, inducing an artificial difference. Overall, our results highlight the necessity for open data sharing efforts such as Abide to provide a more solid ground for the discovery of neuroimaging biomarkers, within the context of the wide human neuroanatomical diversity.

## INTRODUCTION

Autism Spectrum Disorders (ASD) are pervasive developmental disorders with qualitative impairments in social interaction and communication, along with restricted, repetitive, and stereotyped patterns of behaviour. Several cognitive studies have suggested that difficulty integrating multiple sources of stimulation may be a common characteristic of ASD, which has lead for example to the influential “Weak central coherence” hypothesis (1). The neural basis of these difficulties have been hypothesised to be an imbalance between local and distant connections: a local over-connectivity combined with a long-distance under-connectivity (2, 3). The connectivity hypothesis has been a major subject of study and discussion in ASD research (4, 5).

The corpus callosum – the largest commissure connecting the left and right hemispheres of the brain – appeared then as a natural candidate to look for evidence of connectivity abnormalities. The corpus callosum exists exclusively within eutherian mammals (kangaroos and other marsupials lack a corpus callosum), and it has been suggested that it plays an important role in the evolution of functional lateralisation. The number of callosal axons appears to be proportionally smaller in mammals with large brains (like humans) compared with mammals with small brains (like mice). It has been proposed that a proportionally smaller number of callosal fibres, and their increased length, could hinder the formation of interhemispheric synchronous neuronal populations, thus facilitating local recruitment, and possibly leading to the lateralisation of function (language being a classic example) (6–8). The corpus callosum has been in consequence one of the most studied white matter tracts in ASD. Numerous reports have indeed described a statistically significantly smaller CC among ASD patients compared with controls, and a series of studies have suggested a higher incidence of ASD within cases of CC agenesis or callosotomy patients (9, 10).

However, many of these analyses have relied on small samples sizes (in the order of 30 patients versus 30 controls), not having enough statistical power to find medium effect sizes of 0.5 standard deviation between groups. Despite this lack of statistical power, studies often report statistically significant differences. A solution to the methodological problems associated with small sample sizes has been recently proposed by the “Autism Brain Imaging Data Exchange” project, Abide (11). Large sample sizes are difficult to gather and analyse by any single research group. The Abide project provides open access to behavioural data and neuroimaging data (anatomical and functional) for almost 600 patients and 600 age, sex and IQ matched controls from an international consortium of 17 research groups. In addition to providing the research community with the statistical power necessary to detect even small differences in case/control designs, this open dataset allows us also to use more sophisticated analysis strategies and have a wider perspective on neuroanatomical diversity within ASD patients and controls.

In this article we first present a review of previous studies of size differences in the corpus callosum between ASD patients and controls. We observed a general lack of statistical power (only 20% power to detect 2-sided differences of 0.3 standard deviations at 0.05 level of significance), which contrasted with the frequent report of significant findings (12 out of 17 studies). Next, we present our analysis of the diversity of the corpus callosum in the Abide cohort, differences among scanning sites, differences related to age, sex brain size polymorphism and diagnostic group. Even though previous studies have reported diagnostic group differences as large as 0.3-0.7 standard deviations, and despite having analysed a number of subjects comparable to the sum of all previously studied, we did not find any significant difference between ASD patients and controls. Finally, we discuss possible ways in which analysis strategies such as the normalisation of corpus callosum size by total volume, or the matching of subjects by IQ scores, could lead to artefactual differences in brain volume and corpus callosum size between ASD patients and controls.

## METHODS

### 1. Meta-analysis

We included all studies from the recent review on corpus callosum size by Frazier et al (12), and we searched PubMed (http://www.ncbi.nlm.nih.gov/pubmed/) for additional human neuroimaging studies reporting differences in corpus callosum volume between ASD population and a healthy population using the terms: (autism OR PDD OR “pervasive developmental disorder”) AND “corpus callosum”.

From these studies we excluded those that did not report measurements of corpus callosum size and standard deviation for patients and controls. We did not exclude studies because of demographic (for example, gender), clinical (for example, low or high functioning) or methodological characteristics (for example, number of corpus callosum segments analysed).

Studies have used different approaches to the segmentation of the corpus callosum, either measuring it as a single object, or dividing it into 3, 5 or 7 segments. For the sake of comparison, when a global measure of corpus callosum size was not available we combined the mean and standard deviation values for each segment to produce a single estimate. For a corpus callosum segmented into k parts, the mean size was calculated as

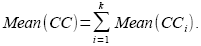

We estimated the global standard deviation of the corpus callosum by assuming (based on our own data) that the size of the region’s segments were correlated with ρ =0.5, and used the formula:

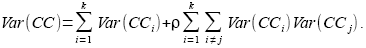

The standard deviation σ is then

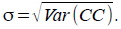

For each article we computed the effect size as a Cohen’s d:

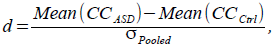

where σ_*Pooled*_ is the pooled standard deviation of ASD and Control subjects computed as:

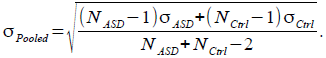

We used G*Power (v 3.1, http://www.psycho.uni-duesseldorf.de/abteilungen/aap/gpower3) to compute for each article the achieved statistical power (for a 2-sided t-test), and their power to detect *a priori* a relatively small effect size of 0.3 standard deviation (2-sided). We used the R (http://www.r-project.org) package Meta to estimate global effect size, heterogeneity and publication bias.

### 2. Analysis of the Abide project cohort

We analysed the size of the corpus callosum in the Abide cohort (http://fcon_1000.projects.nitrc.org/indi/abide). We processed the 1102 subjects with T1-weighted MRI data available using FreeSurfer v5.1. Statistical analyses were performed using JMP Pro 10.0.2 (http://www.jmp.com), R, iPython (http://ipython.org) and G*Power.

MRI data in Abide is labelled with 20 different "Site" levels, corresponding to 17 different scanning sites. Indeed, some sites contributed with more than 1 subject sample (for example, UCLA_1 and UCLA_2, or UM_1 and UM_2). In all cases, however, subjects were scanned using the same scanner and the same parameters, so we combined these site labels into a single one (for example, UCLA or UM).

We developed an open online tool to visually control the accuracy of the segmentations (http://siphonophore.org/qccc). Based on this quality control we excluded 380 subjects. Most of them (N=331) were excluded because the middle segment of the corpus callosum was incorrectly labelled, and included parts of the fornix (Fig. 1). A total of 722 subjects passed the quality control.

**Figure 1.**
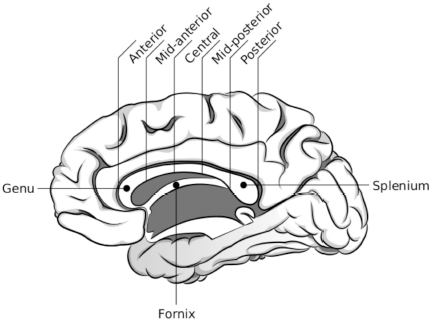
Segmentation of the corpus callosum. The corpus callosum was segmented into 5 regions, from genu to splenium: anterior, mid-anterior, central, mid-posterior and posterior.

The age range of these subjects was from 6.4 years to 64 years. We only included the 694 subjects with ages from 7.5 to 40 years. Tissue segmentation may be unreliable in younger subjects (N=10) because of the ongoing myelination, and brain anatomy may start showing signs of ageing in the older subjects (N=19). The mean age of the retained subsample was 16.8±7.1 years, without statistically significant differences in age distribution between ASD patients and controls (P=0.80, Kolmogorov-Smirnov asymptotic test).

Among the 694 subjects included in our analyses, 328 were ASD patients (290 males, 38 females) and 366 were controls (304 males, 62 females). This sample size provided us 99% power to detect 2-sided differences larger than Cohen’s d=0.3. Because the number of females was statistically significantly larger among the control group (P=0.051, Fisher’s exact test, 2-sided), we also performed our analyses using only males (N=594; 290 patients, 304 controls). Using this male-only sub-sample, we had 95.4% power to detect 2-sided differences larger than Cohen’s d=0.3.

Intelligence Quotients (IQ) were available for 672 of these subjects. Full IQ (FIQ) was available for 590 subjects, and Verbal IQ (VIQ) and Performance IQ (PIQ) were available for 538 subjects. Therefore, there were 82 subjects for whom we had VIQ and PIQ data, but not FIQ data. For those 82 subjects we used a linear model to estimate FIQ based on VIQ and PIQ. This model used data from the 705 subjects that had FIQ, VIQ and PIQ in the complete original cohort. The resulting model – which explained 98% of the variance of FIQ – was:

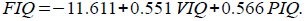

Table 1 provides a decomposition per scanning site of the group of subjects we retained for analysis.

**Table 1.** Description per scanning site of the subjects from the Abide project retained for analysis.

| Site | Institution | N<sub>ASD</sub>/Total | N<sub>ctrl</sub>/Total | Age<sub>ASD</sub>(range) | Age<sub>ctrl</sub>(range) |
| --- | --- | --- | --- | --- | --- |
| Caltech | Caltech Institute of Technology, USA | 7/19 | 8/19 | 26.3(20-39) | 25.2(20-39) |
| CMU | Carnegie Mellon University, USA | 6/14 | 5/13 | 27.5(22-31) | 31.2(25-40) |
| KKI | Kennedy Krieger Institute, USA | 20/22 | 32/33 | 9.9(8-13) | 10.2(8-13) |
| Leuven | University of Leuven, Belgium | 27/29 | 28/35 | 17.4(12-29) | 17.7(12-28) |
| MaxMun | Ludwig Maximilians University Munich, Germany | 11/24 | 22/33 | 21.2(8-35) | 25.4(10-35) |
| NYU | New York University Langone Medical Center, USA | 54/79 | 75/105 | 14.7(8-39) | 16.3(8-32) |
| OHSU | Oregon Health and Science University, USA | 11/13 | 11/15 | 11.1(8-14) | 9.9(8-12) |
| Olin | Olin, Institute of Living at Hartford Hospital, USA | 16/20 | 14/16 | 17.4(12-24) | 16.9(10-23) |
| Pitt | University of Pittsburg, School of Medicine, USA | 30/30 | 27/27 | 18.9(9-35) | 18.9(9-33) |
| SBL | Social Brain Lab, BCN NIC UMC Groningen and Netherlands Institute for Neurosciences, Netherlands | 7/15 | 8/15 | 30.7(27-35) | 32.9(26-39) |
| SDSU | San Diego State University, USA | 8/14 | 12/22 | 15.2(12-17) | 14.3(12-17) |
| Stanford | Stanford University, USA | 8/20 | 5/20 | 9.6(8-12) | 11.2(8-12) |
| Trinity | Trinity Centre for Health Sciences, Ireland | 9/24 | 11/25 | 18.2(14-23) | 17.6(12-26) |
| UCLA | University of California, Los Angeles, USA | 32/62 | 35/47 | 13.3(8-18) | 12.9(10-18) |
| UM | University of Michigan, USA | 23/68 | 27/77 | 12.8(9-19) | 14.8(9-29) |
| USM | University of Utah, School of Medicine, USA | 42/58 | 26/43 | 22.0(11-38) | 22.9(9-39) |
| Yale | Yale Child Study Center, USA | 17/28 | 20/28 | 13.1(9-18) | 12.9(8-18) |
| **Total** |  | **328** | **366** | **16.6(8-39)** | **17.0(8-40)** |

### 3. Analysis of neuroanatomical diversity: allometric scaling

We used allometric scaling to study the way in which the size of the corpus callosum changes relative to total brain volume. The allometric scaling relationship between 2 variables X and Y is formulated as

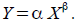

It results from the solution of a growth model where fractional changes in Y are proportional to fractional changes in X: dY/Y=dX/X. Taking logarithms, the allometric scaling relationship can be written as:

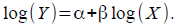

The value β can be then estimated as the slope of the regression line of log(Y) on log(X). If β =1 then Y changes proportionally to X (isometry). On the contrary, if β <1 then the proportion of Y/X decreases as X increases (negative allometry), and if β >1 then the proportion Y/X increases as X increases (positive allometry).

## RESULTS

### A. Meta-analysis

On October 8th, 2013, our PubMed query returned 183 articles, among which 17 fulfilled our inclusion criteria. Combined, these articles provided data on a total of 980 subjects, 521 subjects diagnosed with ASD and 459 controls. Only 77 subjects were females. The mean age was of 14.8 years. Out of the 17 studies, 5 included low-functioning (LF) and high-functioning (HF) subjects, 2 out of 17 included LF only, and 8 out of 17 included HF subjects only. The size of the corpus callosum correlates with total brain volume, and 5 out of 17 studies used the strategy of normalising (divide) CC size by total brain volume, whereas 9 out of 17 used brain volume as a covariate in a linear model. Finally, 8 out of 17 studies matched the ASD and control groups by IQ (Table 2).

**Table 2.** Characteristics of the population represented by the studies reviewed. Matching strategy “Relative”: Differences in brain volume were accounted by dividing CC by BV. Matching strategy “GLM”: Differences in brain volume were accounted by including BV as a covariate in a general linear model. Matching strategy “IQ Match”: Differences in IQ were accounted for by matching groups by intelligence scores.

| Reference | Age ASD/Ctrl | IQ level (LF/HF) | N<sub>ASD</sub>(Females) | N<sub>Ctrl</sub>(Females) | Matching strategy |  |  |
| --- | --- | --- | --- | --- | --- | --- | --- |
|  |  |  |  |  | Relative | GLM | IQ Match |
| 1 Gaffney 1987 (34) | 11±5/12±5 | LF/HF | 13(3) | 35 (14) | No | No | No |
| 2 Egaas 1995 (35) | 16±10/16±10 | LF/HF | 51(6) | 51(6) | No | No | No |
| 3 Piven 1996 (36) | 18±5/20±4 | LF/HF | 35(6) | 36(16) | No | Yes | No |
| 4 Elia 1999 (37) | 11±4/11±3 | LF | 22(0) | 11(0) | No | No | No |
| 5 Manes 1999 (38) | 14±7/12±5 | LF | 27(5) | 17(6) | Yes | No | Yes |
| 6 Rice 2005 (39) | 12±4/13±4 | HF/LF | 12() | 8() | No | Yes | No |
| 7 Vidal 2006 (40) | 10±3/11±3 | HF | 24(0) | 26(0) | Yes | Yes | Yes |
| 8 Boger 2006 (41) | 4±0.3/4±0.5 | – | 45(7) | 26(8) | Yes | Yes | No |
| 9 Alexander 2007 (42) | 16±7/16±6 | HF | 43(–) | 34(–) | No | Yes | Yes |
| 10 Just 2007 (3) | 27±12/25±10 | HF | 18(1) | 18(3) | Yes | No | Yes |
| 11 Hardan 2009 (43) | 11±1/11±1 | HF | 22(0) | 23(0) | Yes | No | Yes |
| 12 Freitag 2009 (44) | 18±4/19±1 | HF | 15(2) | 15(2) | No | Yes | Yes |
| 13 Keary 2009 (45) | 20±10/19±9 | HF | 32(2) | 34(2) | No | Yes | Yes |
| 14 Anderson 2011 (46) | 22±7/21±7 | HF/LF | 53(0) | 39(0) | No | No | Yes |
| 15 Hong 2011 (47) | 9±2/10±2 | HF | 18(0) | 16(0) | No | Yes | Yes |
| 16 Frazier 2012 (48) | 11(8-12)/11(7-13) | HF | 23(0) | 23() | No | Yes | No |
| 17 Prigge 2013 (49) | 14±8/15±7 | HF/HF | 68(0) | 47(0) | No | Yes | Yes |

Table 3 summarises the mean sizes and standard deviation of the CC in the ASD and control groups. The different values were scaled to provided measurements in cm^2^ (This scaling does not affect our meta-analysis, which was performed on standardised mean differences). The standardised effect sizes (mean differences) were not found to be heterogeneous (τ^2^ <0.0001, H=1, I^2^=0%, Q=9.96), and the fixed and random effect meta-analyses provided the same estimates of the mean effect size, ES=-0.51 (95% confidence interval=[−0.63,−0.38], z=−7.642, P<0.0001, Fig. 2).

**Table 3.** Size of the corpus callosum in the studies reviewed, group differences and statistical power. SD: Standard Deviation. P-value in bold are <0.05 (uncorrected). The units from the different studies were scaled to cm2 (provided for reference, as our meta-analysis was performed on standardised values).

|  | Study | Mean<sub>ASD</sub>±SD<br>(cm<sup>2</sup>) | Mean<sub>CTRL</sub>±SD<br>(cm<sup>2</sup>) | Pooled SD<br>(cm<sup>2</sup>) | Effect<br>Size | P-Value<br>(2-sided) | Achieved<br>power<br>(2-sided) | Power to<br>detect SD=0.3<br>(2-sided) |
| --- | --- | --- | --- | --- | --- | --- | --- | --- |
| 1 | Gaffney 1987 (34) | 5.89±1.04 | 6.24±1.37 | 1.31 | -0.27 | 0.070 | 14.6% | 17.3% |
| 2 | Egaas 1995 (35) | 5.57±0.99 | 5.89±0.91 | 0.95 | -0.33 | **0.0011** | 38.2% | 32.3% |
| 3 | Piven 1996 (36) | 6.15±0.83 | 6.40±0.38 | 0.64 | -0.39 | **0.0017** | 36.9% | 24.1% |
| 4 | Elia 1999 (37) | 5.26±1.00 | 5.41±0.64 | 0.91 | -0.16 | 0.36 | 6.4% | 12.7% |
| 5 | Manes 1999 (38) | 4.64±0.99 | 5.71±0.97 | 0.99 | -1.07 | **<0.0001** | 93.6% | 16.2% |
| 6 | Rice 2005 (39) | 7.34±1.11 | 7.75±1.14 | 1.15 | -0.35 | 0.13 | 11.4% | 9.3% |
| 7 | Vidal 2006 (40) | 6.06±1.15 | 6.68±0.79 | 0.99 | -0.62 | **<0.0001** | 57.7% | 17.9% |
| 8 | Boger 2006 (41) | 4.59±0.67 | 4.99±0.72 | 0.70 | -0.57 | **<0.0001** | 67.6% | 24.1% |
| 9 | Alexander 2007 (42) | 7.87±1.99 | 9.32±1.70 | 1.88 | -0.77 | **<0.0001** | 91.4% | 25.2% |
| 10 | Just 2007 (3) | 6.40±0.88 | 7.1±0.88 | 0.89 | -0.78 | **<0.0001** | 62.8% | 13.9% |
| 11 | Hardan 2009 (43) | 5.74±1.13 | 6.58±1.04 | 1.10 | -0.76 | **<0.0001** | 69.8% | 16.2% |
| 12 | Freitag 2009 (44) | 6.22±0.45 | 6.54±1.24 | 0.96 | -0.34 | 0.071 | 14.6% | 12.2% |
| 13 | Keary 2009 (45) | 6.19±1.09 | 6.76±1.10 | 1.11 | -0.51 | **<0.0001** | 54.0% | 22.4% |
| 14 | Anderson 2011 (46) | 6.54±1.20 | 7.05±0.90 | 1.09 | -0.46 | **<0.0001** | 59.4% | 29.6% |
| 15 | Hong 2011 (47) | 8.14±1.31 | 8.27±1.27 | 1.31 | -0.10 | 0.58 | 4.6% | 13.3% |
| 16 | Frazier 2012 (48) | 6.30±1.11 | 6.78±1.08 | 1.11 | -0.43 | **0.005** | 29.9% | 16.7% |
| 17 | Prigge 2013 (49) | 5.74±0.91 | 6.24±0.89 | 0.91 | -0.55 | **<0.0001** | 83.7% | 36.0% |

**Figure 2.**
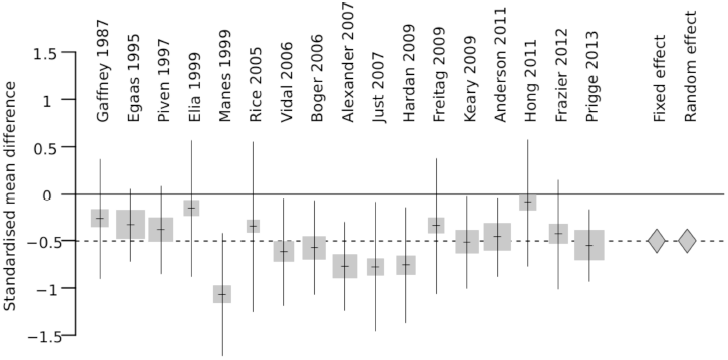
Standardised mean difference in CC size between ASD patients and controls in the articles analysed. Our metaanalysis showed a mean difference of 0.5 standard deviations, smaller in the ASD groups. For each study, the size of the square corresponds with the sample size. The standardised mean difference is shown by diamonds (the estimation was the same for the fixed and random effect models).

A classic test for publication bias estimates the presence of a correlation between the effect size and the standard error of each study (13). A correlation could be detected if small studies were more likely to be published when the effect size is large and then more likely to produce a statistically significant finding. Large studies, which approximate better the true effect size, should then provide a reference. The funnel plot in Fig. 3 appears to be symmetric, and we did not find a significant correlation between the effect size and the standard error of each study (t=0.169, d.f.=15, P-value=0.8681). However, it is difficult to establish whether there was or not a publication bias because there are not really large studies in our sample: the largest statistical power to detect a 0.3 standard deviation was 36%. In general, the mean achieved power was of just 46.9% and the studies appeared to be very underpowered, with only a 20% average power to detect a difference of 0.3 standard deviations (commonly recognised as a “small” difference). Despite the general lack of power, 12 out of 17 studies reported significant results.

**Figure 3.**
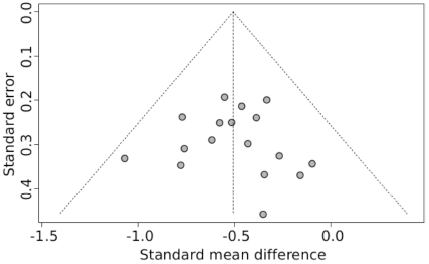
Standard error versus standard mean difference. The funnel plot does not show a bias among articles with small sample sizes (large standard errors) to over-estimate the standard mean difference. However, the absence of studies with sufficiently large cohorts does not allow us to conclude of an absence of publication bias. The dashed lines indicate the 95% confidence interval.

### B. Analysis of the Abide cohort

We start by providing a brief description of the autistic traits and intelligence scores in the ASD group compared with the control group (subsection 1). Next, in subsections 2-4 we describe the differences related to scanning site, the effect of age, sex, and the differences between diagnostic groups in intracranial volume, brain volume, and corpus callosum volume. In subsection 5 we describe the allometric scaling relationship between corpus callosum volume and intracranial volume, and between corpus callosum volume and brain volume. In subsection 6 we describe the correlation between intelligence scores and brain volume. Finally, in subsection 7 we summarise all our previous results to look at the effect that different analysis strategies may have on the detection of differences related to diagnostic group.

#### 1. Characteristics of the Abide population

Several ASD scales were available for different subsets of subjects in the Abide cohort. Scores in the Autism Diagnostic Observation Schedule (ADOS) were available for 435 subjects. The mean score among ASD subjects was 11.9±3.8 (N=383) versus 1.3±1.4 among controls (N=32). The difference was highly significant (t=-34.28, P<0.0001). In the Social Responsiveness Scale the mean score among ASD subjects was 91.3±30.5 (N=190) versus 21.6±16.6 (N=175) among controls. The difference was highly significant (t=-27.44, P<0.0001). Finally, a smaller subset of subjects had also Autism Quotient scores. The mean AQ score among ASD subjects was 30.5±8.1 (N=28) versus 13.3±5.5 among controls (N=28). The difference was again highly significant (t=-9.3, P<0.0001).

Because of the emphasis on functional connectivity analyses in Abide, only high-functioning subjects were studied (it would be more difficult to ask low-functioning subjects not to move, and sedation affects functional imaging results). The mean FIQ was 105±17 among ASD patients and 111±12 among controls, statistically significantly higher in the control group (P<0.0001, 2-sided t-test, t=5.22). There was also a statistically significant difference in mean VIQ: 103±18 in the ASD group and 111±13 in the control group (P<0.001, 2-sided t-test, t=5.61), but there was no statistically significant difference in mean PIQ: 106±17 in the ASD group and 107±13 in the control group (P=0.19, 2-sided t-test, t=1.31). For all 3 IQ values the standard deviation was statistically significantly larger in the ASD group than in the control group (2-sided F-tests for FIQ: P<0.0001, F(318,352)=1.95; VIQ: P<0.0001, F(263,273)=2.10; PIQ: P<0.0001, F(263,273)=1.68). Table 4 shows these differences within the individual scanning sites (note that the minimum sample size required to detect with 80% power a 2-sided difference in IQ larger than 0.5 standard deviations is of N=128, only achieved by the NYU site).

**Table 4.** IQ differences between ASD and Ctrl per scanning site in Abide (negative values mean smaller in the ASD group). (*) None of the controls retained from SBL had IQ scores available. (**) VIQ and PIQ scores were not available. P-values are for 2-sided t-tests (uncorrected).

| Site | N | FIQ | | VIQ | | PIQ | | $\rho$ (VIQ,PIQ) | | |
| --- | --- | --- | --- | --- | --- | --- | --- | --- | --- | --- |
| | | $\Delta$ | P-val | $\Delta$ | P-val | $\Delta$ | P-val | R<sup>2</sup> | $\rho$ | P-value |
| Caltech | 15 | -10.9 | 0.05 | -10.5 | 0.17 | -6.11 | 0.27 | 0.059 | 0.24 | 0.38 |
| KKI (**) | 52 | -16.1 | **0.0008** | - | - | - | - | - | - | - |
| Leuven | 55 | -13.4 | **0.0007** | -19.0 | **0.0001** | -4.7 | 0.24 | 0.099 | 0.32 | **0.019** |
| MaxMun (**) | 33 | -4.8 | 0.34 | - | - | - | - | - | - | - |
| NYU | 129 | -1.9 | 0.46 | -3.7 | 0.15 | 1.3 | 0.65 | 0.19 | 0.43 | **<0.0001** |
| OHSU (**) | 21 | -12.9 | 0.11 | - | - | - | - | - | - | - |
| Olin (**) | 28 | -0.6 | 0.92 | - | - | - | - | - | - | - |
| Pitt | 57 | -0.2 | 0.95 | -0.5 | 0.89 | 0.4 | 0.89 | 0.22 | 0.47 | **0.0003** |
| SBL (*) | 7 | - | - | - | - | - | - | 0.070 | 0.26 | 0.57 |
| SDSU | 20 | 11.5 | 0.16 | 11.6 | 0.14 | 7.9 | 0.30 | 0.43 | 0.64 | **0.0018** |
| Stanford | 13 | 2.6 | 0.72 | -3.0 | 0.72 | 6.4 | 0.43 | 0.11 | 0.33 | 0.27 |
| Trinity | 20 | -8.1 | 0.28 | -8.5 | 0.22 | -6.5 | 0.40 | 0.51 | 0.71 | **0.0004** |
| UCLA | 67 | -5.8 | 0.07 | -5.8 | 0.08 | -3.9 | 0.25 | 0.10 | 0.32 | **0.0082** |
| UM | 50 | -0.3 | 0.95 | -4.3 | 0.44 | 3.7 | 0.38 | 0.03 | 0.18 | 0.22 |
| USM | 68 | -15.3 | **<0.0001** | -18.7 | **<0.0001** | -8.5 | **0.027** | 0.22 | 0.46 | **<0.0001** |
| Yale | 37 | -10.5 | 0.089 | -11.7 | 0.086 | -10.4 | 0.077 | 0.56 | 0.75 | **<0.0001** |

VIQ and PIQ were correlated at ρ =0.44 (we will use these values later). Table 4 shows the strength of this correlation at the different scanning sites (note that the minimum sample size required to detect with 80% power a correlation of 0.3 is of N=67, achieved only by the NYU, UCLA and USM scanning sites).

#### 2. Intracranial volume

Freesurfer estimates ICV from the inverse of the determinant of the linear transformation matrix used to align each individual’s brain to the MNI152 template (see Buckner et al (14) for more details on this method, and a comparison with manual segmentation). This provides an estimation of each individual’s ICV as a proportion of the template’s ICV.

##### Site effect

The mean ICV was 1368±231 cm^3^, with statistically significant differences among centres (P<0.0001, F=32.8, d.f.=16), varying from 232 cm^3^ below the global mean (Stanford) to 273 cm^3^ above the global mean (Caltech). Site effects accounted for 43.7% of ICV variance.

##### Age effect

There was a small but significant increase in ICV of 4.3 cm^3^ per year (P=0.0005, F=12.19, d.f.=1), which accounted for a 1.7% of ICV variance.

##### Sex effect

We observed the expected difference in mean ICV between males and females, 9.4% smaller in females (P<0.0001, F=28.2, d.f.=1). The Sex effect explained 3.9% of total ICV variance.

##### Diagnostic group effect

The mean ICV, adjusted for Site, Age and Sex effects, was of 1331±168 cm^3^. We had 90% power to detect 2-sided differences larger than 41 cm^3^ (a 3.1% difference). ICV was 6.8 cm^3^ larger in the ASD group, but this difference was not statistically significant (GLM for ICV with site, age, sex and diagnostic group as fixed effects, Diagnostic group effect P=0.61, F=0.26, d.f.=1). The complete model explained 46.9% of ICV variance.

#### 3. Brain volume

Supratentorial brain volume was estimated using Freesurfer 5.1, and includes everything except the cerebellum and the brain stem. In particular, it includes the volume of the ventricles, choroid plexus, and vessels.

##### Site effect

The mean BV was 1131±130 cm^3^, with statistically significant differences among centres (P<0.0001, F=10.6, d.f.=16), varying from 267 cm^3^ below the global mean (Stanford) to 157 cm^3^ above the global mean (Trinity). Site effects accounted for 20.1% of BV variance.

##### Age effect

BV showed a small but statistically significant increase of 2.3 cm^3^ per year (P=0.001, F=10.86, d.f.=1), which explained 1.5% of BV variance.

##### Sex effect

BV was statistically significantly smaller among females by 9.3% (P<0.0001, F=62.11, d.f.=1). The Sex effect explained 8.2% of BV variance.

##### Diagnostic group effect

The mean BV, adjusted for site, age and sex effects, was of 1097±112 cm^3^. We had 90% power to detect 2-sided differences larger than 28 cm^3^ (i.e., 2.6% of the average BV). BV was 4.0 cm^3^ larger in the ASD group, but this difference was not statistically significant (GLM for BV with site, age, sex and diagnostic group as fixed effects, diagnostic group effect P=0.86, F=0.03, d.f.=1). The complete model explained 26.0% of BV variance.

BV correlated relatively strongly with ICV, ρ =0.64 (41.5% of the variance, P<0.0001, F=491.6, both BV and ICV adjusted for site, age and sex effects).

#### 4. Corpus callosum

We used the default settings of Freesurfer 5.1, which segments the corpus callosum as a 5 mm thick slab, and divides it into 5 segments of equal length in the main CC axis (closely corresponding with the antero-posterior axis, see Fig. 1).

##### Site effect

The global mean volume of the CC was 3.16±0.54 cm^3^, with statistically significant differences among sites (P<0.0001, F=10.7, d.f.=16). Mean CC volume ranged from 0.61 cm^3^ below the global mean (Stanford), to 0.4 cm^3^ above the global mean (Trinity). Site differences accounted for 20.2% of total CC variance. Mean sizes per subregion, and variance due to site effects are listed in Table 5.

**Table 5.** Mean sizes per callosal subregion, and variance due to site, age, sex and diagnostic group effects in Abide. Corpus callosum slabs were 5 mm thick. The full model includes site, age, sex and diagnostic group as effects. P-value in bold are <0.05 (uncorrected).

|  | Callosal region |  |  |  |  |
| --- | --- | --- | --- | --- | --- |
|  | Posterior | Mid-posterior | Central | Mid-anterior | Anterior |
| **Mean size (cm<sup>3</sup>)** | 0.89±0.18 | 0.44±0.09 | 0.45±0.09 | 0.49±0.13 | 0.90±0.18 |
| **Site effect** |  |  |  |  |  |
| F | 5.63 | 7.81 | 9.19 | 12.87 | 7.74 |
| P-value | **<0.0001** | **<0.0001** | **<0.0001** | **<0.0001** | **<0.0001** |
| R<sup>2</sup> | 11.7% | 15.6% | 17.8% | 23.3% | 15.5% |
| **Age effect** |  |  |  |  |  |
| Increase (mm<sup>3</sup>/year) | 7 | 3 | 2 | 1 | 5 |
| F | 63.30 | 41.54 | 17.07 | 2.99 | 31.66 |
| P-value | **<0.0001** | **<0.0001** | **<0.0001** | 0.084 | **<0.0001** |
| R<sup>2</sup> | 8.4% | 5.7% | 2.4% | 0.4% | 4.4% |
| **Sex effect** |  |  |  |  |  |
| Percent difference (1-Female/Male) | 8.4% | 6.8% | 4.5% | 7.6% | 8.2% |
| F | 16.12 | 8.74 | 4.54 | 7.28 | 15.17 |
| P-value | **<0.0001** | **0.0032** | **0.033** | **0.0072** | **<0.0001** |
| R<sup>2</sup> | 2.3% | 1.2% | 0.7% | 1% | 2.1% |
| **Group effect** |  |  |  |  |  |
| Difference (mm<sup>3</sup>) | -4.7 | 1.3 | -3.3 | -1.4 | 1.7 |
| F | 0.63 | 0.01 | 0.67 | 0.14 | 0.04 |
| P-value | 0.43 | 0.91 | 0.41 | 0.71 | 0.84 |
| R<sup>2</sup> | 0.00 | 0.00 | 0.00 | 0.00 | 0.00 |
| **Variance explained by the full model** | 15.1% | 16.3% | 18.3% | 24.7% | 17.1% |

##### Age effect

There was a small but statistically significant increase in mean CC size of 19 mm^3^ per year (P<0.0001, F=45.0, d.f.=1), which accounted for 6.1% of CC variance. The increase was more pronounced in the Posterior and Anterior subregions, which are those that more strongly correlate with BV (see below). Table 5 lists Age effects for the 5 callosal subregions, and the amount of variance they account for.

##### Sex effect

Mean CC size was statistically significantly smaller among females by 7.4% (P<0.0001, F=17.1, d.f.=1) compared with males. The Sex effect explained 2.4% of CC variance. A similar difference was observed in all subcallosal regions, especially the Posterior and Anterior ones (Table 5). We will discuss in the next section the extent to which this difference is explained by the significant difference in BV between females and males.

##### Diagnostic group effect

The mean CC volume adjusted for Site, Age and Sex effects was of 3.08±0.48 cm^3^. We had 90% power to detect differences larger than 119 mm^3^ (a 3.9% difference). The observed difference was < 7 mm^3^ (larger in controls), not statistically significant (P=0.56, F=0.35,d.f.=1), as were none of the differences in the 5 callosal subregions (Table 5). The full model for total CC volume explained 22.0% of the variance.

The different callosal subregions presented a medium correlation among them (Table 6). The 1st principal component of the variance/covariance matrix of callosal subregion volume explained 65% of the variance, and was correlated with ρ =0.43 with BV and with ρ =0.98 with total CC volume. This principal component corresponded to a coordinated increase in the size of the posterior and anterior subregions. Indeed, the posterior and anterior subregions of the corpus callosum showed one of the strongest correlations among them: ρ =0.61.

**Table 6.** Pearson’s correlation matrix among ICV, BV and callosal regions.

|  | ICV | BV | Posterior | Mid-posterior | Central | Mid-anterior | Anterior |
| --- | --- | --- | --- | --- | --- | --- | --- |
| ICV | 1 | 0.67 | 0.26 | 0.12 | 0.13 | 0.11 | 0.33 |
| BV | 0.67 | 1 | 0.34 | 0.30 | 0.33 | 0.29 | 0.40 |
| Posterior | 0.26 | 0.34 | 1 | 0.58 | 0.47 | 0.39 | 0.61 |
| Mid-posterior | 0.12 | 0.30 | 0.58 | 1 | 0.67 | 0.57 | 0.48 |
| Central | 0.13 | 0.33 | 0.47 | 0.67 | 1 | 0.69 | 0.46 |
| Mid-anterior | 0.11 | 0.29 | 0.39 | 0.57 | 0.69 | 1 | 0.36 |
| Anterior | 0.33 | 0.40 | 0.61 | 0.48 | 0.46 | 0.36 | 1 |

#### 5. Allometric scaling of the corpus callosum with ICV and BV

The total corpus callosum volume presented a small correlation with ICV (ρ =0.27) and a medium correlation with BV (ρ =0.43). The same was observed for individual callosal regions, that showed a small correlation with ICV and medium correlation with BV (Table 6). A partial correlation analysis suggested that the variability of the CC was more directly related to BV than to ICV, and that the observed correlation between ICV and CCV was largely mediated by BV (Table 7). In consequence, our further analyses related CC variability with BV.

**Table 7.** Matrix of partial correlations among ICV, BV and CC. Most of the correlation between ICV and CC was mediated by the correlation between ICV and BV.

|  | ICV | BV | CC |
| --- | --- | --- | --- |
| ICV | – | 0.64 | -0.027 |
| BV | 0.64 | – | 0.35 |
| CC | -0.027 | 0.35 | – |

Our results show a statistically significant negative allometry of CC with BV, with a scaling coefficient β =0.64. We observed the same negative scaling for the callosal regions (average β =0.64, range from 0.58 to 0.73, Table 8).

**Table 8.** Allometric scaling coefficient (β) of the callosal regions and ICV and BV. P-value in bold are <0.05 (uncorrected).

| Region | ICV |  |  |  | BV |  |  |  |
| --- | --- | --- | --- | --- | --- | --- | --- | --- |
| | $\beta$ | R<sup>2</sup> | F | P | $\beta$ | R<sup>2</sup> | F | P |
| Posterior | 0.34 | 5.0% | 36.7 | **<0.0001** | 0.59 | 9.1% | 69.7 | **<0.0001** |
| Mid-posterior | 0.19 | 1.4% | 10.5 | **0.0013** | 0.58 | 8.4% | 64.2 | **<0.0001** |
| Central | 0.20 | 1.9% | 13.4 | **0.0003** | 0.59 | 10.9% | 84.2 | **<0.0001** |
| Mid-anterior | 0.24 | 1.8% | 12.6 | **0.0004** | 0.71 | 9.7% | 74.5 | **<0.0001** |
| Anterior | 0.47 | 9.9% | 75.9 | **<0.0001** | 0.73 | 15.0% | 122.3 | **<0.0001** |
| CC | 0.32 | 6.9% | 51.4 | **<0.0001** | 0.64 | 17.8% | 149.6 | **<0.0001** |

The scaling coefficient for CC was not different between ASD patients and controls – neither the diagnostic group effect nor the interaction between diagnostic group and BV were statistically significant (Diagnostic group P=0.43, F=0.62, d.f.=1; Diagnostic group * BV P=0.85, F=0.04, d.f.=1). Additionally, none of the callosal regions showed any significant diagnostic group effect or diagnostic group * BV effect.

Because of the allometric relationship between CC and BV, and due to the significant effect of sex on BV, we expect females to have smaller CC than males in absolute terms (as we observed in the previous section), but larger than males in relative terms (i.e., CC/BV). This was indeed the case: relative CC was statistically significantly larger in females than males (Age and Site as covariates, Sex effect: F=6.75, P=0.0095, d.f.=1). The difference was, however, completely explained by the relationship between CC and BV, as adding BV as a covariate made the sex effect not statistically significant (Age, Site and BV as covariates, Sex effect: F=1.8, P=0.18, d.f.=1) as it had been previously pointed out by Luders et al (15).

#### 6. Correlation between brain volume and IQ

IQ is often used to match ASD patient groups and controls. But several studies have reported a significant correlation between IQ and ICV or IQ and BV. Additionally, IQ appears to be very sensitive to environmental factors such as stress or socio-economic status. It is then not clear that the relationship between IQ and brain size would be the same in a control group and a group of ASD patients. Indeed, whereas we did observe the expected correlation between IQ and BV in the control group (ρ =0.23, P<0.0001, F=18.75), the relationship was significantly weaker in the ASD patients group (ρ =0.04, P=0.044, F=4.10). A GLM for FIQ with Site, Age and Sex as covariates, and BV, diagnostic group and diagnostic group * BV as main effects, indicated a statistically significant diagnostic group effect (P<0.0001, F=29.33, d.f.=1), as well as a statistically significant interaction between BV and diagnostic group (P=0.0178, F=5.64, d.f.=1). In other words, whereas in controls FIQ increased on average by 1 point every 31 cm^3^, the same 1 point increase in FIQ required an increase of 84 cm^3^ in ASD patients.

Most of the diagnostic group * BV interaction effect on FIQ appeared to be due to a difference in the correlation between VIQ and BV in ASD patients and controls, but not between PIQ and BV. VIQ and BV correlated with ρ =0.22 in controls, but with ρ =0.08 in ASD patients. By contrast, PIQ and BV correlated with ρ =0.18 and ρ =0.17 in controls and patients, respectively (see Table 9 for correlations at individual sites). A GLM for VIQ with Site, Age and Sex as confounding covariates and BV, diagnostic group and diagnostic group * BV as main effects indicated that the interaction effect was statistically significant (P=0.035, F=4.5, d.f.=1). This was not the case for the interaction effect diagnostic group * BV on PIQ (P=0.60, F=0.27, d.f.=1).

**Table 9.** Correlation between FIQ, VIQ, PIQ and BV. P-value in bold are <0.05 (uncorrected).

| Site | N | FIQ |  |  | VIQ |  |  | PIQ |  |  |
| --- | --- | --- | --- | --- | --- | --- | --- | --- | --- | --- |
| | | R<sup>2</sup> | $\rho$ | P | R<sup>2</sup> | $\rho$ | P | R<sup>2</sup> | $\rho$ | P |
| Caltech | 15 | 0.062 | -0.25 | 0.37 | 0.002 | 0.04 | 0.87 | **0.015** | -0.12 | 0.66 |
| CMU | - | - | - | - | - | - | - | - | - | - |
| KKI | 52 | 0.001 | -0.03 | 0.84 | - | - | - | - | - | - |
| Leuven | 55 | 0.080 | 0.28 | **0.036** | 0.077 | 0.28 | **0.040** | **0.021** | 0.15 | 0.29 |
| MAX | 33 | 0.001 | 0.03 | 0.84 | - | - | - | - | - | - |
| NYU | 129 | 0.070 | 0.26 | **0.0024** | 0.030 | 0.17 | **0.049** | 0.070 | 0.26 | **0.0025** |
| OHSU | 21 | 0.005 | -0.07 | 0.77 | - | - | - | - | - | - |
| Olin | 28 | 0.007 | 0.08 | 0.68 | - | - | - | - | - | - |
| PITT | 57 | 0.023 | 0.15 | 0.26 | 0.009 | 0.09 | 0.48 | **0.012** | 0.11 | 0.42 |
| SBL | 7 | 0.004 | -0.06 | 0.89 | 0.017 | 0.13 | 0.78 | 0.192 | -0.44 | 0.32 |
| SDSU | 20 | 0.006 | 0.08 | 0.7378 | 0.064 | 0.25 | 0.28 | **0.010** | -0.09 | 0.70 |
| Stanford | 13 | 0.014 | 0.12 | 0.6905 | 0.047 | 0.22 | 0.48 | 0.0 | 0.0 | 0.96 |
| Trinity | 20 | 0.02 | 0.14 | 0.4852 | 0.020 | 0.14 | 0.54 | **0.033** | 0.18 | 0.44 |
| UCLA | 67 | 0.035 | 0.19 | 0.1280 | 0.012 | 0.11 | 0.38 | 0.050 | 0.22 | 0.068 |
| UM | 50 | 0.173 | 0.42 | **0.0026** | 0.03 | 0.17 | 0.21 | 0.249 | 0.50 | **0.0002** |
| USM | 68 | 0.051 | 0.23 | 0.0635 | 0.010 | 0.10 | 0.40 | 0.078 | 0.28 | **0.021** |
| Yale | 37 | 0.014 | -0.12 | 0.4795 | 0.000 | 0.0 | 0.92 | **0.045** | -0.21 | 0.21 |

#### 7. Impact of the relationship between BV, ICV, CC and IQ in controls and ASD patients of the Abide cohort on analysis strategies.

Our analyses suggested that BV could be used as a parameter indexing a relatively large part of the neuroanatomical diversity in ICV and CC in our cohort, as well as smaller but statistically significant part of the diversity in IQ scores. Summarising from our previous analyses:

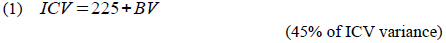

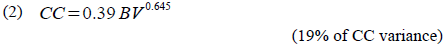

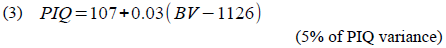

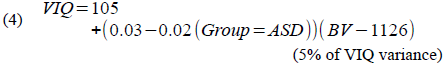

From Equations 3, 4 and the linear model for FIQ in the Abide cohort described in the Methods section we can deduce an expression for the dependance of FIQ on BV:

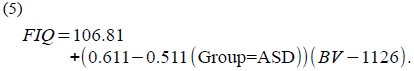

Since the first observations by Kanner (16), many researchers have reported a higher incidence of macrocephaly (17) in ASD, as well as a larger average brain volume (18). Recent studies, however, have shed some doubts on this hypothesis (19, 20). In any case, if head circumference or brain volume were on average larger among ASD patients compared with the general population, and because of Eq. 2, we would expect the proportion of the corpus callosum to be smaller in the ASD group than in the control group. The size of the corpus callosum is not directly proportional to BV, but more like the – power of BV (i.e., to brain “surface” – the surface of the envelope of the brain, not to be confounded with cortical surface).

Many previous studies use either corpus callosum size relative to brain volume (5 out of 17 in our meta-analysis), or include brain volume as a covariate in a linear model (8 out of 17 in our meta-analysis). But if the relationship between CC and BV is non-linear as stated by Eq. 2, and if the groups differed in average brain volume, both procedures could potentially produce artefactual group effects (this would be in general the case for any other comparison of two populations with different average brain volumes – males and females, or tall and short persons, for example).

Human brain volumes are widely variable, with large brains reaching up to 2 times the volume of small brains. If relative CC size were used (i.e., CC/BV) instead of absolute CC, we should expect the proportion of the CC in the largest brains to be up to 1.28-fold (i.e., 2^1-0.644^) smaller than in the smallest brains. Were the brains of ASD patients larger on average than those of the control population, a proportionally smaller CC would be expected.

We used simulations to test the extent to which BV normalisation controlled for differences in BV between populations. We simulated two populations of 50, 100, …, 350 subjects per group, that had a standardised mean BV difference of 0.1, 0.2, …, 1.0 standard deviations. We generated BV values distributed as in the Abide data, and corresponding CC values based on Eq. 2. As in Abide, 19% of the variance of CC was explained by BV. The result is summarised in Figure 4. We observed that BV normalisation did not eliminate the effect of mean BV differences between groups when this difference was large enough (depending on group size). For example, BV normalisation in a study based on two groups of 50 subjects each with a mean BV difference of 0.65 standard deviations will detect a statistically significant difference in CC size 50% of the time, i.e., a statistical power of 50%. For a statistical power of 20% (the average of the studies in our meta-analysis), a mean BV difference of 0.25 standard deviations will suffice.

**Figure 4.**
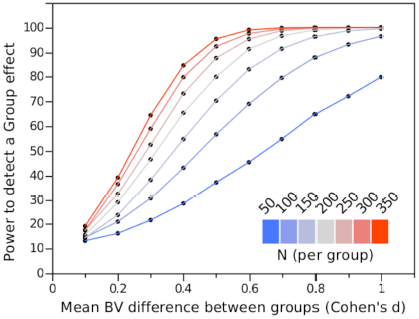
Statistical power to detect a significant, artefactual, difference in CC size for two groups presenting a difference in mean BV: CC normalised by BV. Statistical power as a function of the difference in mean BV between 2 simulated groups (Cohen’s d) consisting of 50, 100, … 350 subjects each. CC is normalised by dividing it by BV. Because of the non-linear relationship between CC and BV, CC normalisation was not sufficient to control for the difference in mean BV.

Alternatively, if brain volume (or intracranial volume) were used as a covariate in a linear model, the non-linear Eq. 2 would be approximated by a linear function. The expansion of Eq. 2 in Taylor series around a reference brain volume μ is:

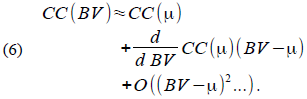

A linear approximation of Eq. 2 results from discarding factors related to the 2^nd^ derivative and higher, and will be valid close to μ. We obtain

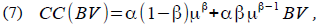

were α =0.39 and β =0.64. If we were to study only 1 group, the best value of μ would be close to the average brain volume of the group. But if we studied two groups with different average brain volumes, a linear model with a group effect and an interaction effect would allow for 2 intercepts and slopes in Eq. 7. Given enough statistical power, these difference may become significant.

We used simulations to test the extent to which including BV and BV*group effects controlled for differences in BV between populations. As before, we simulated two populations of 50 to 350 subjects per group, with standardised mean BV differences of 0.1 to 1 standard deviations, and CC values following Eq. 2 (19% of its variance). The result is summarised in Figure 5. This approach was better than the precedent at controlling the effect of mean BV differences between groups, and induced detectable differences in CC size only for very large groups and very large differences in mean BV.

**Figure 5.**
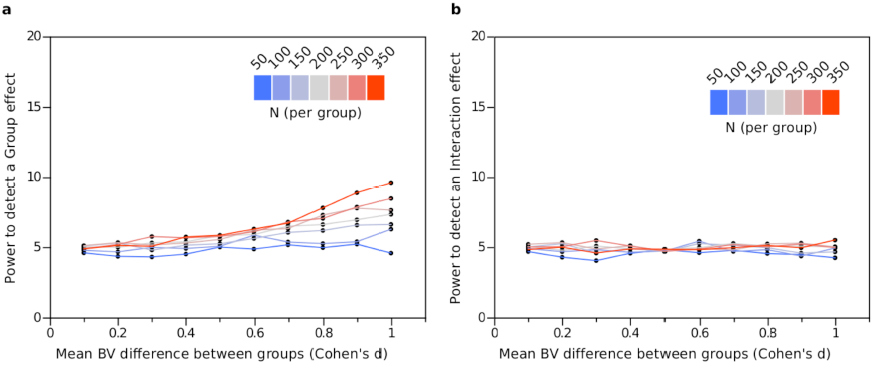
Statistical power to detect a significant, artifactual, difference in CC size between two groups presenting a difference in mean BV: BV used as covariate in a GLM. (a) Statistical power to detect a group effect in CC size as a function of the difference in mean BV between 2 simulated groups (Cohen’s d) consisting of 50, 100, …, 350 subjects each, (b) Statistical power to detect an interaction effect BV*Group in CC size as a function of the difference in mean BV between 2 simulated groups consisting of 50, 100, …, 350 subjects each. Including BV as a covariate successfully controls for BV effects even for large sample sizes and large differences in mean BV.

Another frequent strategy in the study of ASD neuroanatomy has been to match patients and controls by IQ (used by 9 out of 17 studies in our meta-analysis). Equations 4 and 5 suggest, however, that the relationship between IQ and brain size may be different in patients and controls. If larger increases in brain volume are required among patients than among controls to obtain a similar increase in IQ, matching groups by IQ may bias the recruitment to either decrease the number of controls with large brain volumes, or increase the number of patients with large brain volumes. This bias could in turn affect the assessment of group differences in CC size.

We evaluated through simulation the impact of matching by FIQ two groups with different correlations between FIQ and BV. We considered, based on the Abide data, two groups with the same mean BV and with the same correlation between PIQ and BV:

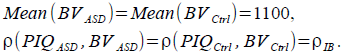

In one group (the simulated ASD group), VIQ did not correlate with BV and was also lower than in the control group (Ctrl):

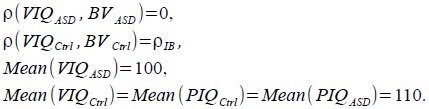

In all cases VIQ and PIQ correlated at 0.5, the standard deviation of VIQ and PIQ was of 15 points, and the standard deviation of BV was of 100 cm^3^:

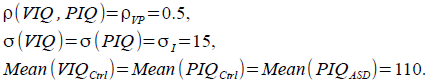

Then, the variance matrices between VIQ, PIQ, BV for ASD (Σ_*ASD*_) and Ctrl (Σ_*Ctrl*_) groups were, respectively:

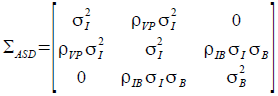

and

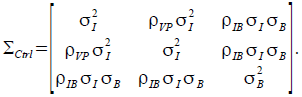

The only difference between this variance-covariance matrices is the correlation between VIQ and BV, which is zero in the ASD group and ρ_*IB*_ σ_*I*_ σ_*B*_ in the Ctrl group.

Next, we computed FIQ scores for each subject as

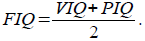

To match the groups we matched their FIQ histograms. We sorted all subjects by FIQ and retained alternatively one ASD subject followed by one Ctrl subject. As expected, matching by FIQ eliminated more often ASD subjects than controls in the small BV end, and more controls than ASD subjects in the large BV end. We repeated this procedure for groups with 50, 150, …, 750 subjects, and correlations ρ_*IB*_ between IQ and BV equal to 0.3, 0.35, …, 0.5. For each group size and ρ_*IB*_ value we performed 1000 simulations.

We observed that based on the strength of the correlation between IQ scores and BV, matching by FIQ induced an increasing difference in mean BV between groups. The relationship was linear (Fig. 6a):

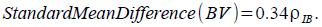

We counted for each group size and ρ_*IB*_ value the number of times that a statistically significant mean BV difference was found (Fig.6b). Only in groups with a strong relationship between IQ and BV the strategy of matching groups by FIQ induced a reproducible difference in mean BV. For example, comparisons between 2 groups of 50 subjects each, with a correlation between IQ and BV or ρ_*IB*_ = 0.5 will detect a significant difference in mean BV 20% of the time (which is, sadly, higher than the average statistical power of many neuroimaging studies (21)).

**Figure 6.**
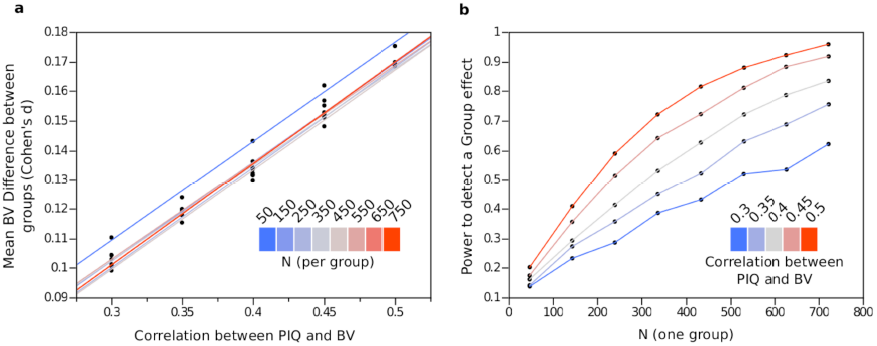
Effect of IQ matching on mean BV. Two groups were simulated with different correlation between FIQ and BV. In one (the control group) both VIQ and PIQ correlated with BV. In the other (the ASD group) only PIQ correlated with BV. FIQ was computed as the average between VIQ and PIQ, and subjects in both groups were selected to match the FIQ distributions, (a) Mean BV difference induced by matching two populations as a function of the correlation between PIQ and BV, for groups of 50, 150, …, 750 subjects each, (b) Power to detect a difference in mean BV induced by matching two populations by FIQ as a function of the number of subjects per group. Statistical power curves were drawn for correlations between PIQ and BV of 0.3, 0.35,…,0.5.

Finally, because the size of callosal subregions is highly determined by total CC size, group differences will be more easily detected in the posterior and anterior subregions, which are more strongly correlated with total CC size, and with total BV. Indeed, the 1st principal component of the variance-covariance matrix of CC subregion size explains 65% of the variance and correlates at ρ =0.98 with CC size:

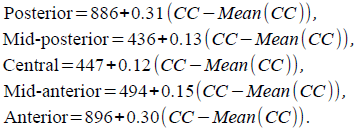

## DISCUSSION AND CONCLUSION

Our analyses of the neuroanatomical variability of the corpus callosum and the brain volume in the Abide cohort did not show any statistically significant difference between patients and controls, contrary to the majority of the previous reports. The difference that we observed between ASD patients and controls was < 0.02 standard deviations for the mean size of the corpus callosum, and < 0.04 standard deviations for the brain volume, in other words, practically inexistent.

This absence of difference did not seem to be due to a difference in the characteristics of the Abide cohort. The Abide cohort is comparable with the previous ones in regards to the subject’s age, sex ratio and IQ level. Additionally, nothing indicates that we may be dealing with an ASD population presenting a milder autistic phenotype – the ADOS, AQ and SRS scores were all highly significantly different between patients and controls. Several of the previous reports, however, included low-functioning subjects either exclusively or in addition to high-functioning individuals, whereas the Abide cohort includes only high-functioning subjects. But differences in corpus callosum size had been equally reported among low-functionning and high-functionning subjects. Out of the 9 studies in our meta-analysis that included exclusively high-functioning subjects, 7 reported significant differences in corpus callosum size as large as 0.78 standard deviations.

The absence of difference does not seem to stem either from the multi-centric nature of Abide. There are several scanning sites within Abide that included as much or even more subjects than in the previous literature (NYU: N=129, USM: N=68, UCLA: N=67). In none of them we detected a statistically significant difference (Table 10 summarises the significance of the group effects for ICV, BV, CC, and CC segments for all scanning sites. The number of uncorrected significant P-values was the expected given the number of multiple tests). Finally, the computational neuroanatomy methods that we used to segment and measure CC, BV and ICV were all standard and well validated. We are making available our data tables and scripts as well as a web interface with our quality control decisions to allow the community to inspect and criticise our analyses.

**Table 10.** Summary of diagnostic group effects in BV, ICV and CC per scanning site in Abide. P-value in bold are <0.05 (uncorrected).

| Site | BV | ICV | CC | Posterior | Mid-posterior | Central | Mid-anterior | Anterior |
| --- | --- | --- | --- | --- | --- | --- | --- | --- |
| Caltech | 0.19 | 0.43 | 0.82 | 0.92 | 0.65 | 0.94 | 0.66 | 0.84 |
| CMU | 0.57 | 0.16 | 0.97 | 0.69 | 0.41 | 0.98 | 0.59 | 0.82 |
| KKI | 0.41 | 0.20 | **0.024** | 0.13 | 0.094 | **0.012** | 0.059 | 0.36 |
| Leuven | 0.27 | 0.69 | 0.24 | 0.21 | 0.76 | 0.26 | 0.66 | 0.11 |
| MaxMun | 0.48 | 0.58 | 0.23 | 0.26 | 0.59 | 0.81 | 0.91 | 0.022 |
| NYU | 0.48 | 0.26 | 0.97 | 0.66 | 0.97 | 0.86 | 0.14 | 0.37 |
| OHSU | 0.20 | 0.55 | 0.93 | 0.56 | 0.91 | 0.48 | 0.70 | 0.92 |
| Olin | 0.48 | 0.49 | 0.19 | 0.64 | **0.040** | 0.85 | 0.74 | **0.044** |
| Pitt | 0.52 | 0.27 | 0.25 | 0.52 | **0.035** | 0.17 | **0.050** | 0.39 |
| SBL | 0.20 | 0.49 | **0.038** | 0.12 | 0.17 | 0.71 | 0.99 | **0.025** |
| SDSU | 0.41 | 0.089 | 0.89 | 0.60 | 0.57 | 0.85 | 0.58 | 0.94 |
| Stanford | 0.38 | 0.89 | 0.85 | 0.75 | 0.51 | 0.70 | 0.73 | 0.76 |
| Trinity | 0.86 | 0.80 | 0.27 | 0.32 | 0.11 | 0.18 | 0.68 | 0.51 |
| UCLA | 0.86 | 0.39 | 0.16 | 0.75 | 0.20 | **0.020** | **0.040** | 0.47 |
| UM | 0.30 | 0.46 | 0.62 | 0.73 | 0.10 | 0.80 | 0.080 | 0.98 |
| USM | 0.58 | 0.60 | 0.12 | 0.074 | 0.63 | 0.57 | 0.11 | 0.10 |
| Yale | 0.41 | 0.43 | 0.28 | 0.38 | 0.88 | 0.17 | **0.048** | 0.37 |

The major, clear, difference between our analysis and the previous ones was statistical power. Using the Abide cohort we achieved 99% power to detect differences of 0.3 standard deviations, whereas the highest statistical power to detect this effect size among the studies in our meta-analysis was of only 36%. Given the low mean statistical power of the previous reports (20% on average), even if there were a real difference in corpus callosum, there should be about 80% of negative reports. However, only 5 out of the 17 articles analysed reported non-significant differences. It has been observed that the scarcity of negative results is especially marked in autism research (22). This is a very damaging tendency within our field, which impedes us from deciding on which hypothesis are worth pursuing and which are not.

Despite the appeal of the hypothesis of a smaller corpus callosum in ASD, we need to consider the possibility that it may not be true. Does this falsify or weakens the underconnectivity theory? The underconnectivity theory states that autism is caused by “insufficient integration circuitry” (3). But whereas many articles in different sub-fields of autism research have indicated that their findings support (verify) the underconnectivity hypothesis, it is not clear what finding would be necessary to prove it wrong (falsify it). As pointed out by Braitenberg (23), most of the brain tissue could be considered as circuitry (myelinated and non myelinated axons, dendrites), the main role of which is undoubtedly some type of integration. It is not entirely surprising that ASD, as other psychiatric disorders, can be interpreted as some type of insufficiency in integration circuitry. We believe that to progress, autism research requires a theoretical framework with stronger, more clearly falsifiable predictions. In particular, a smaller corpus callosum has not been directly stated as a prediction of the underconnectivity theory. If our aim were to preserve the theory, we could simply add an ad-hoc clause (“autism is caused by an insufficiency in integration circuitry of a type that does not change the size of the corpus callosum”), but the appropriateness of this approach has been criticised (24).

Our results suggest that non-linear variations in corpus callosum size relative to brain volume present in the general population, or different patterns of covariation of confounding factors could lead to some of the group differences reported in the literature. We found that 19% of the variance in CC size was captured by a relationship with brain volume where progressively larger brains had a proportionally smaller CC. If for some reason the brain volume in one population were different than in the other, this difference could lead to a group difference in corpus callosum size (for example, if females and males were compared). Our analyses showed that normalisation of CC size (divide CC by BV, which supposes that the relative size of the CC is independent of BV, i.e., isometric scaling) may not be sufficient to control for a difference in BV. Including BV as a covariate provided a more reliable control, and should be preferred to normalisation (the only more reliable method being, of course, to match groups by brain size as in (15)). Besides a real difference in BV we showed that IQ matching could under certain circumstances induce an artefactual BV difference, that could be later observed as a CC size difference (this will be the case for any other confounding variable that changes differently in cases and controls). Finally, besides the allometric scaling of CC and BV, similar non-linear relationships have been also observed between total cortical surface and BV, between folding (local and global) and BV, and between white matter volume and BV. Because of these nonlinear scaling relationships it is expected that subjects with larger brain volumes will have a larger cortical surface area (25), more folded particularly in the prefrontal cortex (26), and with a larger frontal white matter volume (27). These are exactly the findings that have been reported in several articles comparing ASD patients and controls (2, 28–30). The extent to which they arise from a difference in brain volume (real or artefactual) has yet to be evaluated.

We found that intelligence scores do not covary with BV in the same way in ASD and controls. This is not completely surprising, as FIQ – and VIQ in particular – are known to be affected by environmental factors such as stress or socioeconomic status (31). Psychiatric disorders such as ASD do not only impose to the patients the cognitive challenges that we most often use to define them, but also a daily confrontation with various comorbidities, and various degrees of social, educational and daily-life difficulties depending of our society’s ability to integrate them. This additional burden is very likely to leave physiological traces, in particular neuroanatomical. Finding the biological markers of the causes of ASD may then require to disentangle them not only from risk factors, but also from the social effects of being different (carrying a handicap, being disadvantaged, etc.).

Research suggest today that the aetiology of ASD is highly heterogeneous, with hundreds of genetic mutations associated with it (32, 33), as well as many environmental factors. The patient’s phenotypes are also so diverse, with the presence of such a large number of different comorbidities and wide spectrum of cognitive abilities that one could wonder about the pertinence of trying to look for neuroanatomical traits common to all of them. It is important, however, to remember that the nervous system is strongly self-regulated, and capable of the most striking plasticity. The processes leading to the formation of a mammalian nervous system are incredibly resilient and able to produce viable cognitive function under the most extreme circumstances. The phenomena that we may be able to observe at the scale of the complete nervous system will be more likely the trace of this common mechanism of developmental canalisation and compensation than a direct reflexion of the heterogeneous aetiology. In this sense, the comprehension of the normal response of the nervous system to perturbation, and the way in which the diversity of this response is regulated by the complete individual’s genetic background, may turn to be as important as the direct study of the pathologic cause. The study of large cohorts with extensive behavioural, genetic and neuroimaging data will be of fundamental relevance for this endeavour.

## References

1. Frith U (2003) Autism: Explaining the Enigma (Wiley).

2. Courchesne E, Pierce K (2005) Why the frontal cortex in autism might be talking only to itself: local over-connectivity but long-distance disconnection. Curr Opin Neurobiol 15:225–30.

3. Just MA, Cherkassky VL, Keller TA, Kana RK, Minshew NJ (2007) Functional and Anatomical Cortical Underconnectivity in Autism: Evidence from an fMRI Study of an Executive Function Task and Corpus Callosum Morphometry. Cereb Cortex 17:951–961.

4. Mengotti P, Brambilla P (2014) in Comprehensive Guide to Autism, eds Patel VB, Preedy VR, Martin CR (Springer New York), pp 911–927. Available at: http://link.springer.com/referenceworkentry/10.1007/978-1-4614-4788-7_47 [Accessed January 21, 2014].

5. Hughes JR (2009) Update on autism: A review of 1300 reports published in 2008. Epilepsy Behav 16:569–589.

6. Ringo JL, Doty RW, Demeter S, Simard PY (1994) Time Is of the Essence: A Conjecture that Hemispheric Specialization Arises from Interhemispheric Conduction Delay. Cereb Cortex 4:331–343.

7. Rilling JK, Insel TR (1999) Differential expansion of neural projection systems in primate brain evolution. Neuroreport 10:1453–9.

8. Aboitiz F, López J, Montiel J (2003) Long distance communication in the human brain: timing constraints for inter-hemispheric synchrony and the origin of brain lateralization. Biol Res 36:89–99.

9. Booth R, Wallace GL, Happé F (2011) Connectivity and the corpus callosum in autism spectrum conditions: insights from comparison of autism and callosal agenesis. Prog Brain Res 189:303–317.

10. Paul LK et al. (2007) Agenesis of the corpus callosum: genetic, developmental and functional aspects of connectivity. Nat Rev Neurosci 8:287–299.

11. Di Martino A et al. (2013) The autism brain imaging data exchange: towards a large-scale evaluation of the intrinsic brain architecture in autism. Mol Psychiatry.

12. Frazier TW, Hardan AY (2009) A meta-analysis of the corpus callosum in autism. Biol Psychiatry 66:935–941.

13. Rothstein H, Sutton AJ, Borenstein M (2005) Publication bias in meta-analysis: prevention, assessment and adjustments (Wiley, Chichester, England; Hoboken, NJ).

14. Buckner RL et al. (2004) A unified approach for morphometric and functional data analysis in young, old, and demented adults using automated atlas-based head size normalization: reliability and validation against manual measurement of total intracranial volume. NeuroImage 23:724–38.

15. Luders E, Toga AW, Thompson PM (2014) Why size matters: Differences in brain volume account for apparent sex differences in callosal anatomy: The sexual dimorphism of the corpus callosum. NeuroImage 84:820–824.

16. Kanner L (1943) Autistic disturbances of affective contact. Nervous Child 2:217–250.

17. Lainhart JE et al. (2006) Head Circumference and Height in Autism: A Study by the Collaborative Program of Excellence in Autism. 2274:2257–2274.

18. Courchesne E, Campbell K, Solso S (2011) Brain growth across the life span in autism: Age-specific changes in anatomical pathology. Brain Res 1380:138–145.

19. Chaste P et al. (2013) Adjusting Head Circumference for Covariates in Autism: Clinical Correlates of a Highly Heritable Continuous Trait. Biol Psychiatry 74:576–584.

20. Raznahan A et al. (2013) Compared to what? Early brain overgrowth in autism and the perils of population norms. Biol Psychiatry 74:563–575.

21. Button KS et al. (2013) Power failure: why small sample size undermines the reliability of neuroscience. Nat Rev Neurosci 14:365–376.

22. Joober R, Schmitz N, Annable L, Boksa P (2012) Publication bias: What are the challenges and can they be overcome? J Psychiatry Neurosci JPN 37:149–152.

23. Braitenberg V (2001) Brain Size and Number of Neurons: An Exercise in Synthetic Neuroanatomy. 71–77.

24. Popper KR (2000) The logic of scientific discovery (Routledge, London; New York).

25. Toro R et al. (2008) Brain Size and Folding of the Human Cerebral Cortex. Cereb Cortex 18:2352–2357.

26. Toro R et al. (2008) Brain size and folding of the human cerebral cortex. Cereb Cortex 18:2352–7.

27. Toro R et al. (2009) Brain volumes and Val66Met polymorphism of the BDNF gene: local or global effects? Brain Struct Funct 213:501–9.

28. Carper RA, Moses P, Tigue ZD, Courchesne E (2002) Cerebral lobes in autism: early hyperplasia and abnormal age effects. NeuroImage 16:1038–1051.

29. Herbert MR et al. (2004) Localization of white matter volume increase in autism and developmental language disorder. Ann Neurol 55:530–40.

30. Amaral DG, Schumann CM, Nordahl CW (2008) Neuroanatomy of autism. Trends Neurosci 31:137–45.

31. Hanscombe KB et al. (2012) Socioeconomic Status (SES) and Children’s Intelligence (IQ): In a UK-Representative Sample SES Moderates the Environmental, Not Genetic, Effect on IQ. PLoS ONE 7:e30320.

32. Toro R et al. (2010) Key role for gene dosage and synaptic homeostasis in autism spectrum disorders. Trends Genet TIG 26:363–372.

33. Huguet G, Ey E, Bourgeron T (2013) The genetic landscapes of autism spectrum disorders. Annu Rev Genomics Hum Genet 14:191–213.

34. Gaffney GR, Kuperman S, Tsai LY, Minchin S, Hassanein KM (1987) Midsagittal magnetic resonance imaging of autism. Br J Psychiatry 151:831–833.

35. Egaas B, Courchesne E, Saitoh O (1995) Reduced size of corpus callosum in autism. Arch Neurol 52:794–801.

36. Piven J, Bailey J, Ranson BJ, Arndt S An MRI study of the corpus callosum in autism. Am J Psychiatry 154:1051–1056.

37. Elia M et al. (2000) Clinical Correlates of Brain Morphometric Features of Subjects With Low-Functioning Autistic Disorder. J Child Neurol 15:504–508.

38. Manes F et al. (1999) An MRI study of the corpus callosum and cerebellum in mentally retarded autistic individuals. J Neuropsychiatry Clin Neurosci 11:470–474.

39. Rice SA et al. (2005) Macrocephaly, corpus callosum morphology, and autism. J Child Neurol 20:34–41.

40. Vidal CN et al. (2006) Mapping Corpus Callosum Deficits in Autism: An Index of Aberrant Cortical Connectivity. Biol Psychiatry 60:218–225.

41. Boger-Megiddo I et al. (2006) Corpus Callosum Morphometrics in Young Children with Autism Spectrum Disorder. J Autism Dev Disord 36:733–739.

42. Alexander AL et al. (2007) Diffusion tensor imaging of the corpus callosum in Autism. NeuroImage 34:61–73.

43. Hardan AY et al. (2009) Corpus callosum volume in children with autism. Psychiatry Res Neuroimaging 174:57–61.

44. Freitag CM et al. (2009) Total Brain Volume and Corpus Callosum Size in Medication-Naïve Adolescents and Young Adults with Autism Spectrum Disorder. Biol Psychiatry 66:316–319.

45. Keary CJ et al. (2009) Corpus Callosum Volume and Neurocognition in Autism. J Autism Dev Disord 39:834–841.

46. Anderson JS et al. (2011) Decreased Interhemispheric Functional Connectivity in Autism. Cereb Cortex 21:1134–1146.

47. Hong S et al. (2011) Detecting abnormalities of corpus callosum connectivity in autism using magnetic resonance imaging and diffusion tensor tractography. Psychiatry Res Neuroimaging 194:333–339.

48. Frazier TW, Keshavan MS, Minshew NJ, Hardan AY (2012) A Two-Year Longitudinal MRI Study of the Corpus Callosum in Autism. J Autism Dev Disord 42:2312–2322.

49. Prigge MBD et al. (2013) Corpus Callosum Area in Children and Adults with Autism. Res Autism Spectr Disord 7:221–234.

